# MiR-505-3p is a Repressor of the Puberty Onset in Female Mice miR-505-3p and the puberty onset

**DOI:** 10.1101/402784

**Authors:** Yuxun Zhou, Li Tong, Maochun Wang, Xueying Chang, Sijia Wang, Kai Li, Junhua Xiao

## Abstract

Puberty onset is a complex trait regulated by multiple genetic and environmental factors. In this study, we narrowed a puberty related QTL down to a 1.7 Mb region on chromosome X in female mice and inferred miR-505-3p as the functional gene.

We conducted ectopic expression of miR-505-3p in the hypothalamus of prepubertal female mice through lentivirus-mediated orthotopic injection. The impact of miR-505-3p on female puberty was evaluated by the measurement of pubertal events and histological analysis. The results showed that female mice with overexpression of miR-505-3p in the hypothalamus manifested later puberty onset timing both in vaginal opening and ovary maturation, followed by weaker fertility lying in the longer interval time between mating and delivery, higher abortion rate and smaller litter size. We also constructed miR-505-3p knockout mice by CRISPR/Cas9 technology. MiR-505-3p knockout female mice showed earlier vaginal opening timing, higher serum gonadotrophin and higher expression of puberty-related gene, as well as its target gene *Srsf1* in the hypothalamus than their wild type littermates.

*Srsf1* was proved to be the target gene of miR-505-3p that played the major role in this process. The results of RIP-seq (RNA Immunoprecipitation-sequencing) showed that SF2, the protein product of *Srsf1* gene, mainly bound to ribosome protein (RP) mRNAs in GT1-7 cells. The collective evidence implied that miR-505-3p/SRSF1/RP could play a role in the sexual maturation regulation of mammals.

**Author summary:** The puberty onset in mammals is a vital biological process that signals the acquisition of reproductive capacity. The initiation of puberty is triggered by the activation of hypothalamic pulsatile GnRH surge. The dysregulation of pubertal development shows relevance to later health risks of type 2 diabetes, cardiovascular disease, breast cancer and other health disorders. Recent progress indicates that a lot of genes play a role in the excitatory or inhibitory regulation of GnRH release. However, the detailed pathway of pubertal timing remains unclear. Our previous studies isolated an X-linked QTL that was associated with the timing of puberty in mice. In this study, we proved that miR-505-3p was a female puberty onset regulator based on data from positional cloning, ectopic expression and knockout mouse models. We also assigned *Srsf1* as the functional target gene of miR-505-3p underlying this process. The results of RIP-seq showed that SF2, the protein of *Srsf1* gene, preferential bound to ribosome protein (RP) mRNAs in GT1-7 cells. We propose that miR-505-3p/SF2/RP could play a role in the sexual maturation regulation of mammals.

## Introduction

Puberty onset is an essential and complicated physiology process that relies on the activation of hypothalamic-pituitary-gonadal axis(HPGA).The pulsatile secretion of GnRH hormones from the specialized neurons in the hypothalamus trigger gonadal hormones secreting cascade that results in the maturation of sexual behaviors and signal the transition from non-reproductive juvenile into a fertility competent adult [1, 2]. The dysregulation of pubertal development shows relevance to later health risks for type 2 diabetes, cardiovascular disease, breast cancer and other health disorders [3].

The genetic and environmental factors involved in the regulation of puberty timing have been well characterized. The mutations in Gpr54 (a G protein–coupled receptor) gene could cause autosomal recessive idiopathic hypogonadotropic hypogonadism (iHH) in human, and the puberty regulating function of GPR54 was also revealed in mice with complementary genetic approaches [4–6], which suggested a crucial role of GPR54 and its ligand Kisspeptin in the regulation of puberty. During the last decade, novel central neuroendocrine pathways in the control of puberty have been revealed. In 2017, genome-wide association studies including genotype data of up to ∼370,000 women identified 389 independent signals for age at menarche, ∼250 genes were implicated via coding variation or associated expression[7].The maternal imprinted gene MKRN3 was the first gene identified with an inhibitory effect on GnRH secretion. It was related with central precocious puberty by whole-exome sequencing of pedigree samples [8]. System biology strategies revealed that at least three gene networks might contribute to the puberty onset [9].

In our previous studies, we isolated a QTL (quantitative trait locus) on Chromosome X affecting the vaginal opening in female mice [10]. In this study, We narrowed the QTL down to a 1.7Mb region by constructing 8 interval-specific congenic strains (ISCSs) between C57/BL6 (B6) and C3H/He(C3H) mice. Among the genes in this region, miR-505-3p was assumed to be the potential candidate according to the variation in its flanking sequence and gene expression levels between C3H and B6 mice, as well as its functional annotation. MicroRNAs are endogenous, small non-coding RNAs (∼22nt in length) silencing the gene expression at post-transcriptional level by targeting the 3’ untranslated region (UTRs) of mRNAs. MicroRNAs get involved in various biological processes [11]. Recent evidence suggested that microRNAs also participated in the precise regulation of puberty onset [12]. MiR-505-3p usually serves as a biomarker of various diseases including primary biliary cirrhosis, Parkinson’s disease and inflammatory bowel disease [13–15]. Zucchi et al detected the downregulation of miR-505-3p in newborn rat brain in response to the prenatal stress [16]. Moreover, miR-505-3p was proved to induce apoptosis in MCF7-ADR cells, and functioned as a tumor suppressive miRNA [17], its first validated target by experiments was the alternative splicing factor/splicing factor2 (ASF/SF2 or SRSF1) [18], which is a regulator of mTOR (the mammalian Target of Rapamycin) pathway [19], while the activated mTOR signaling could accelerate the vaginal opening in female rats [20]. Such indirect evidence tempted us to clarify the authentic relationship between miR-505-3p and the mammalian puberty onset regulation.

In this study, we conducted overexpression of miR-505-3p in the hypothalamus of prepubertal female mice through lentivirus-mediated orthotopic injection. We also constructed miR-505-3p knockout mice (miR-505-3p -/-) by CRISPR/Cas9 technology. The impact of miR-505-3p on the female puberty was evaluated by the measurement of pubertal events and histological analysis in these genetically modified mice. The results showed that female mice with ectopic expression of miR-505-3p in the hypothalamus manifested later puberty onset timing, compromised fertility, higher abortion rate and smaller litter size. While miR-505 knockout female mice showed lightly earlier vaginal opening and shorter interval time between mating and delivery. *Srsf1* was proved to be the target gene of miR-505-3p that played the major role in this process. The results of RNA Immunoprecipitation-Sequencing (RIP-Seq) showed SF2, the protein of *Srsf1* gene, mainly bound ribosome protein (RP) mRNAs in GT1-7 cells.

All these results suggest that miR-505-3p may regulate the puberty onset via modulating the expression of *Srsf1* gene and RP. They give us insights into a new regulating pathway involving microRNA, SF2 and ribosome proteins in mammalian puberty onset.

## Results

### MiR-505-3p was identified as a functional candidate gene in the QTL on chromosome X

We have identified a 9.5Mb (2.5cM) QTL that regulates the puberty onset of female mice on chromosome X in our previous work. To narrow down the QTL further, 8 interval-specific congenic strains (ISCSs) with intervals within this region of B6 chromosome X were substituted into C3H background in the N7 generation, in which we recorded the age at VO and body weights of all female mice. Nonparametric tests showed that strain# 1∼3, 7 and 8 showed significantly different VO age from the parental strain C3H, while strain #4∼6 didn’t. Based on the QTL information and allele distribution among those strains, we ascertained a shrunk QTL of about 1.6Mb between rs13483770 and rs299055848 (Fig 1A, S1 Table).

**Fig 1.**
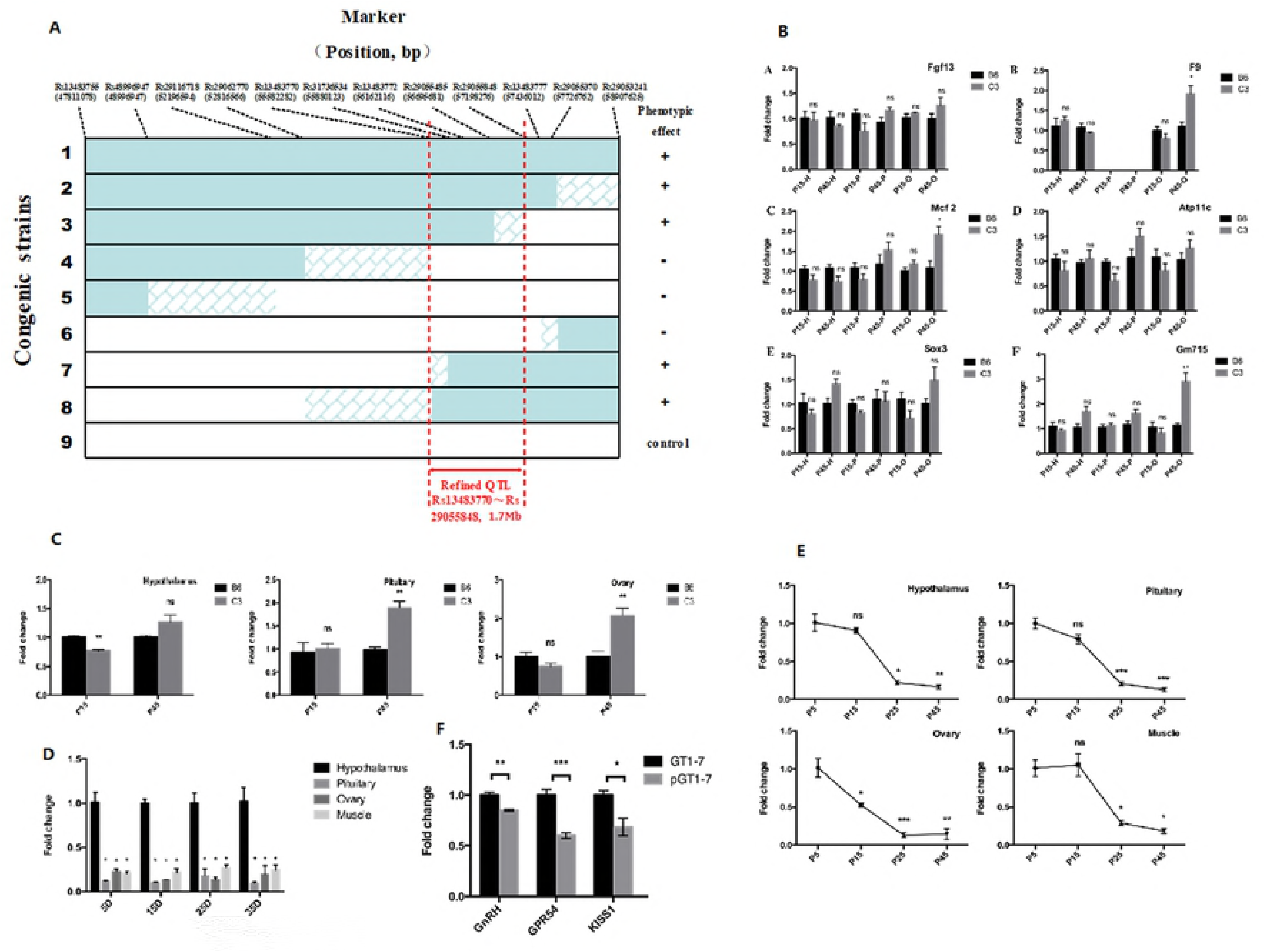
The evidence of miR-505-3p being a candidate gene in the puberty onset related QTL on chromosome X. A: schematic representation of the ISCSs and control strains. The genotype on chromosome X for each of the 8 congenic strains is represented by the horizontal bars. Green portions indicate a known homozygous B6 segment, blank portions represent a known homozygous C3 segment, and hatched regions depict an area where a crossover between C3 and B6 occurred. The dotted lines indicate the boundary of the refined QTL. B: The expression situation of six protein-coding genes in the QTL in HPG axis in B6 and C3 strains. H: hypothalamus; P: pituitary; O:overy. C: The expression difference of miR-505-3p in HPG axis between B6 and C3H mice. D: The Spatial-specific expression of miR-505-3p in different time point of B6 mice. E: The temporal-specific expression of miR-505-3p in different tissues of B6 mice. F: The expression level of puberty related genes in GT1-7 and pGT1-7cells. Bars are means and vertical bars represent SEM (* P<0.05, ** P<0.01, *** P<0.001).

There are 19 genes in the QTL region (rs13483770∼rs299055848) on chromosome X, including seven pseudogenes, four ncRNAs, seven protein-coding genes, and one microRNA gene. To screen for the candidate gene(s), we referred to Sanger database to search for the DNA sequence variations between C3H and B6 mice in this region. Data showed that there were no SNPs, short indels or structural variants lying in the coding region, 5’ or 3’ UTR of these genes except for mir-505, as sensible variations of 5 consecutive SNPs existed near 5’ upstream region of mir-505 gene (S2 Table). To confirm the results from Sanger database, we did DNA sequencing for the interested regions of the 19 genes, and found no exceptions. We compared the mRNA expression levels of the seven protein-coding genes in HPG axis between B6 and C3H female mice in order to find out the differentially expressed genes. Six of them were expressed in mouse HPGA except for the gene of Gm7073, but they made hardly significant differences in the amount of accumulation between the two strains, especially at the hypothalamus or pituitary (Fig 1B). On the other hand, miR-505-3p of B6 was expressed higher in the hypothalamus than that of C3H (Fig 1C) at postnatal 15 days, and lower in the pituitary and ovary at postnatal 45 days. We investigated the temporal and spatial expression pattern of miR-505-3p in B6 female mice, and found that mir-505-3p was most abundant at hypothalamus in the four tested tissues including pituitary, ovary and muscles (Fig1D); it reached its peak value of expression at neonatal period (PD5) and decreased gradually afterwards (Fig1E), which implied its inhibiting role in the sexual maturation process. The integrated evidence suggested that miR-505-3p deserved further investigation as a functional candidate gene.

We constructed a GT1-7 cell line with stable overexpression of miR-505-3p named pGT1-7. In pGT1-7 cells, the expression level of miR-505-3p increased by 1500 times than the GT1-7 cells(S1 Fig). We compared the expression profiling between pGT1-7 and negative control group GT1-7 (transfected with pLenti6.3-nc lentivirus) on DNA microarray, and found that some important puberty related genes, including *Kiss1* and *GnRH*, decreased in pGT1-7 cells. These results were confirmed by qRT-PCR assay (Fig1F), implying that miR-505-3p may participated in the puberty onset regulation through the inhibition of puberty related genes.

### Ectopic expression of miR-505-3p in the hypothalamus influences the puberty onset timing and fertility of female mice

In order to investigate the influence of miR-505-3p on the puberty timing in female mice, we made miR-505-3p overexpressed in the hypothalamus of C57BL/6 (B6) female mice at postnatal 15 days by lentivirus-mediated orthotopic injection. In the window of post injection 5 to 40 days, the hypothalamic miR-505-3p overexpression maintained at a high level, reaching the peak at 10 days after injection (miR-505-3p-LV-treated=24.89±3.319, saline-treated=1.016±0.08129, untreated=1.096±0.2130, P<0.0001). (S2 Fig)

After injection, the mice were weighted once every other day, and the miR-505-3p-LV-treated (abbreviated as LV-treated) female mice showed significant growth retardation compared with the untreated ones (Figure 2A). The vaginal opening in untreated mice were 2 days earlier than the LV-treated ones (P<0.01) (Figure 2B).

**Fig 2.**
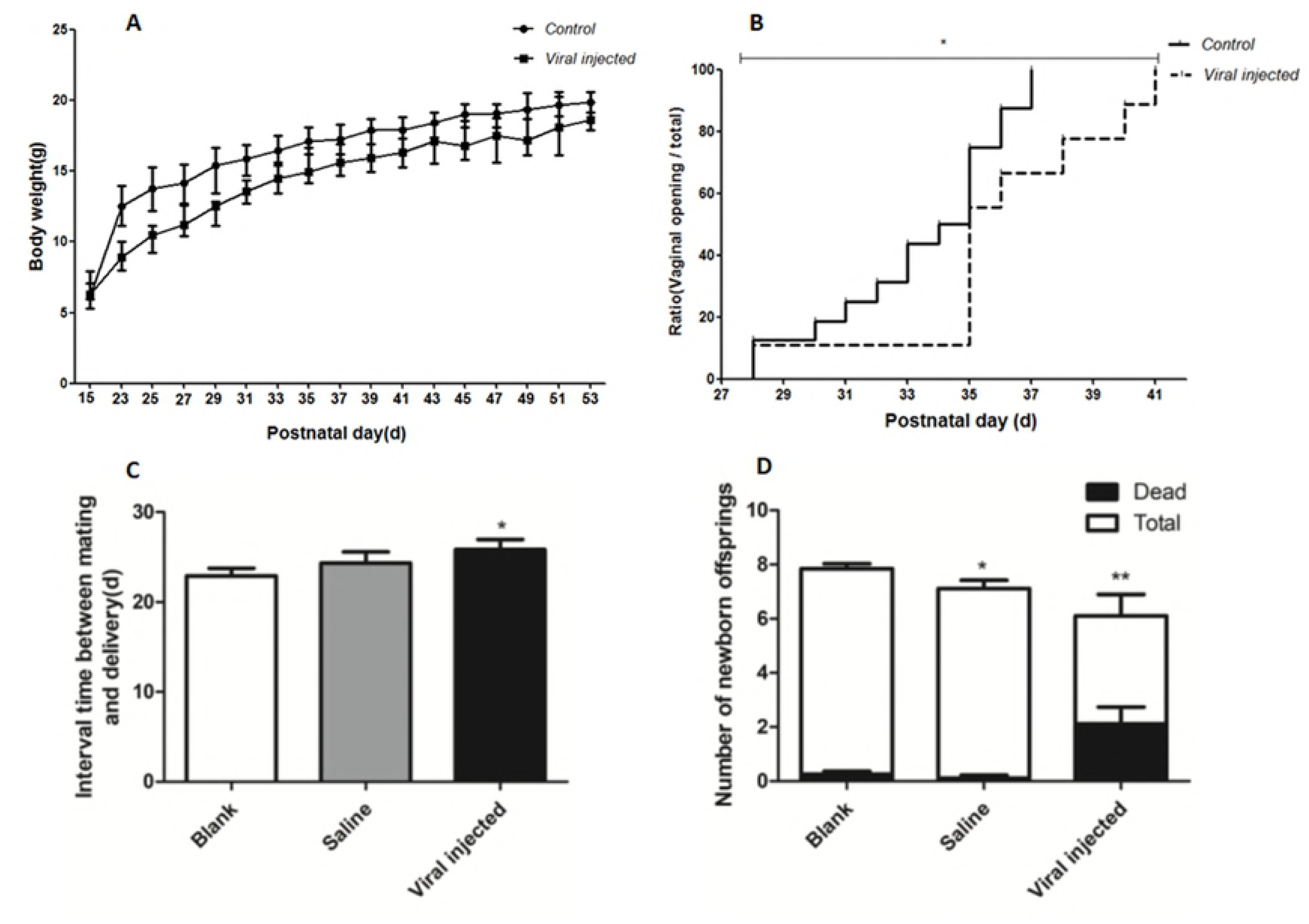
The influence of miR-505-3p ectopic expression in the hypothalamus on the body weight, VO timing and fertility of the tested mice. A: the growth curve of the mice B: the VO time of the tested mice; C: The interval time between mating with male mice and delivery (LV-treated=25.83 ±1.144d, saline-treated=24.33 ±1.247d,untreated=22.92 ±0.8307, P<0.05,*); D: the death rate of offspring before weaning of the tested mice (newborn offspring:LV-treated=6.100 ±0.7944, saline-treated=7.111 ±0.3093, untreated=7.949 ±0.1798,; dead offspring:LV-treated=2.1110±0.6256, saline-treated=0.1111 ±0.1111, untreated=0.2564 ±0.1085. P<0.05 *, P<0.01**)

After sexual maturation, the female mice were mated with wild-type experienced male mice to evaluate the long-term impact of hypothalamic miR-505-3p overexpression on reproduction. The LV-treated female mice needed more time to procreate and had smaller newborn litters compared with the control ones, the death rate of offspring before weaning increased in LV-treated mice as well (Figure2C and 2D). The Hematoxylin-eosin (HE) stained ovarian sections at various time showed the delay of the formation of the preovulatory follicles and corpora luteas (CLs) in LV-treated female mice compared with untreated ones, which provided explanation for the later birth of newborn offspring in LV-treated mice, and the reduced number of newborn offsprings could be attributed to the deficiency or reduced number of preovulatory follicles (Figure 3).

**Fig 3.**
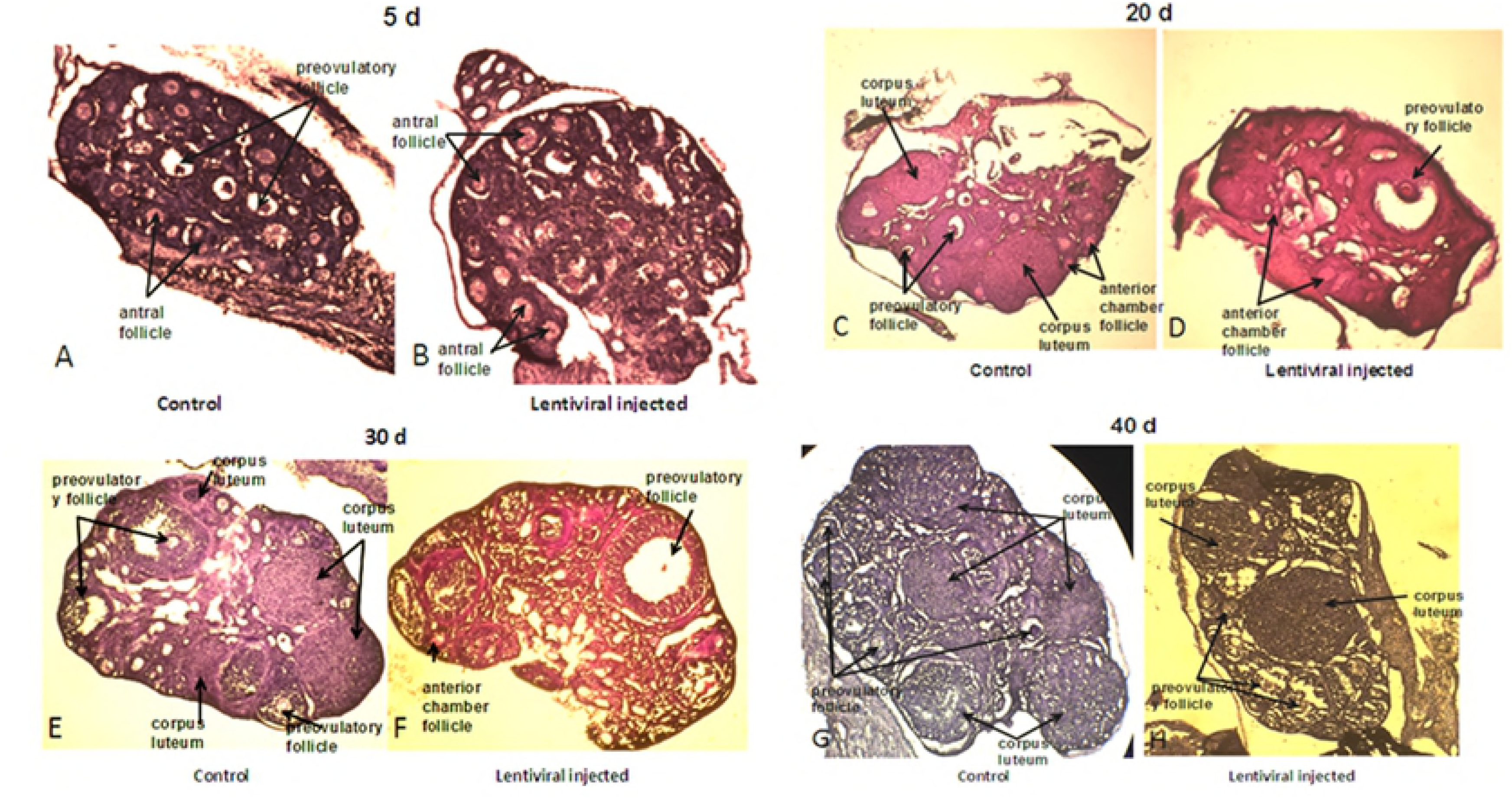
The HE stained ovarian sections at various time

We found higher death rate of litters in LV-treated groups, and the rate of abortion in LV-treated groups was 2.5 -fold higher than the two control groups. About 8 percent of LV-treated mice were sterile in contrast with the control groups, in which no sterile mice were found (Figure 4A). The FISH (florescence in situ hybridization) results of the brain section showed that the sterile female mouse had stronger fluorescence signals near the third ventricle than the signals of its counterparts which had four or eight litters, respectively. This finding suggested the level and the location of miR-505-3p overexpression in the hypothalamus may influence the fertility phenotype of female mice(Figure 4B).

**Fig 4.**
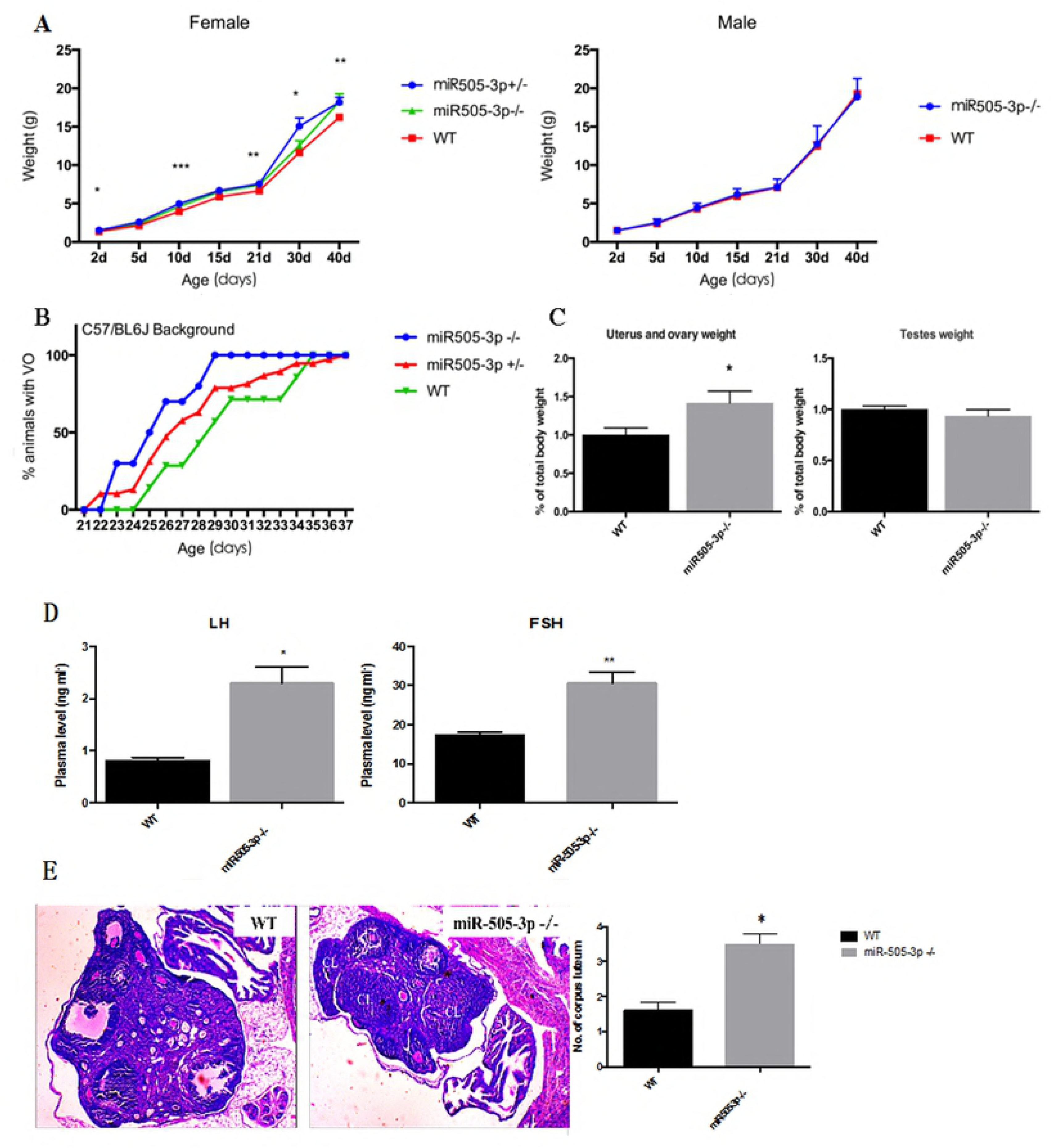
Fertility analysis and in situ hybridization of the brain section. A: Pie chart shows the percentage of female mice which had different number of dead litters and the rate of abortion and infertility among the three groups; B: Images of coronal sections showed the fluorescence signals of hypothalamic miR-505-3p detected by in situ hybridization among one sterile, one reduced fertility, one normal, saline-treated and untreated mice using U6 miR-505-3p and scrambled LNA probes, respectively.

### MiR-505-3p knockout mice showed an advanced onset of puberty and abnormal reproductive phenotypes

With the help of CRISPR/Cas9 technology, we obtained 41 living offsprings, and 16 of them carried mutations in mir-505 cDNA region on their genome detected by DNA sequencing. Two pups with the largest deletion (-17bp and -23bp, respectively) adjacent to the 5’ end of miR-505 (S3 Fig) and in the coding region of mature miR-505-5p were kept as founders to generate DKO miR-505-3p -/-), SKO miR-505-3p +/-) and wild type mice for puberty onset trait investigation by backcrossing to B6 mice. In DKO mice, the expressing level of miR-505-3p could hardly be tested, while in SKO mice, the expressing level of miR-505-3p decreased by approximate half (S4 Fig). In female knockout mice, we observed an increased growth rate and a larger body weight than wild type mice (Fig 5A), 3.57d and 3.87d advance in vaginal opening in SKO and DKO mice (Fig 5B), respectively. At PND45, the mass of reproductive system was larger in knockout mice, indicating advanced sexual development in knockout mice (Fig 5C).

**Fig 5:**
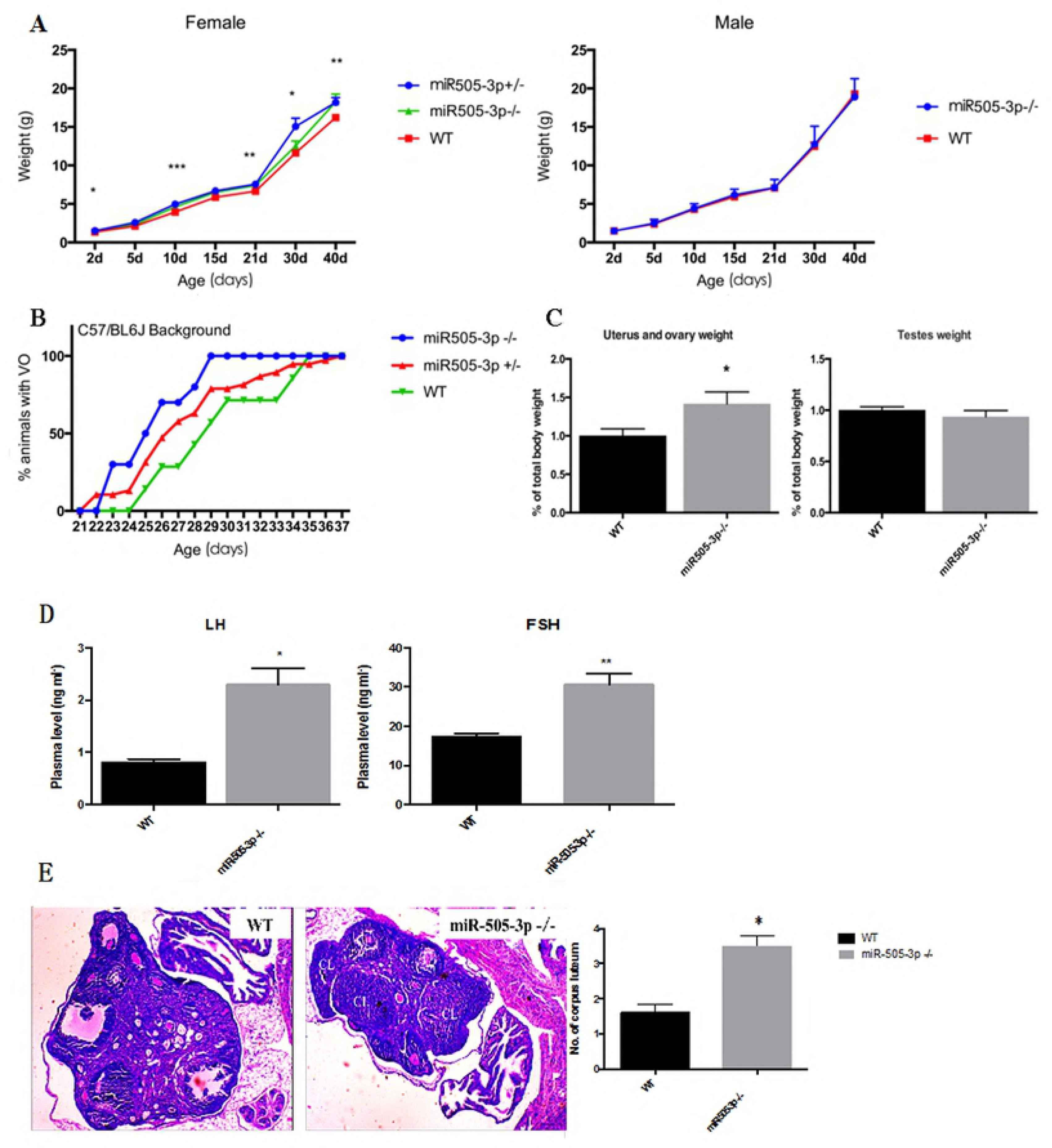
The phenotype of miR-505-3p knockout mice. A: The growth curve of the DKO, SKO and WT mice; B: the percentage of female mice attaining VO at different time point; C: the mass of reproductive system at PND45; D: The serum LH and FSH of the DKO and WT mice; E: HE stained ovarian sections of the tested mice. (*: P<0.05, **: P<0.01)

We then analyzed serum levels of the pituitary gonadotropins luteinizing hormone (LH) and follicle-stimulating hormone (FSH), and the results showed a remarkable increase in female knockout mice (Fig 5D). Moreover, the size of the litter was larger in knockout mice than wild type, and heterozygous knockout mice showed more dystocia in female mice and more dead off springs at 48h (Table 1). Hematoxylin-eosin stained ovarian sections of miR-505-3p knockout mice showed more corpus luteum (Fig 5E). *Srsf1, Kiss1* and *Gnrh* in hypothalamus of miR-505-3p knockout mice and wild mice at different postnatal days were also detected, and the results showed knockout mice had higher expression levels (Fig. 6).

**Table 1.** Effect of miR-505-3p deficiency on female fertility.

| Strains | Litters | Dystocia in female<br>(%) | Mean litter<br>size | $P^b$ | Dead offspring<br>(% at 48h) |
| --- | --- | --- | --- | --- | --- |
| WT | 30 | 0 | 6.032 (0.3485) |  | 1.07 |
| miR-505-3p +/- | 35 | 31.4 | 7.125 (0.3677) | 0.0373 | 29.8 |

Note: a: Values are the mean (s.e.m.). b: P value for mean litter size compared to wild type.

**Fig 6.**
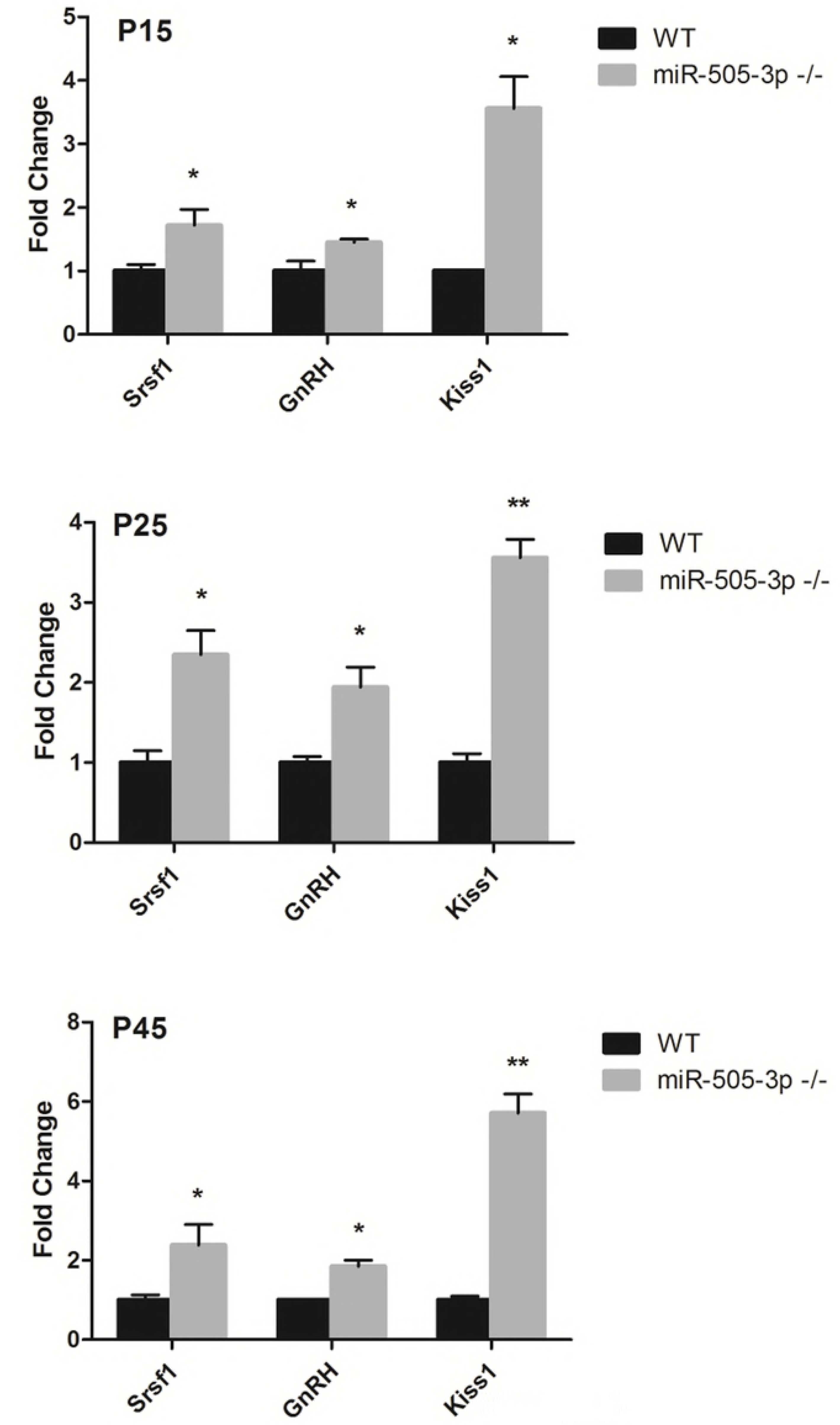
The expression level of Srsf1, Kiss1 and GnRH in the hypothalamus of mice at PND15, PND25, PND45.

### *Srsf1* was proved to be a target gene of miR-505-3p in GT1-7 cell line

We performed the Gene ontology analysis on the gene expression profilings data from the microarray detection between pGT1-7 and GT1-7 cells. 1490 genes were upregulated and 604 genes were downregulated in the pGT1-7 cells. Among these genes, we screened for the miR-505-3p target genes on the TargetScan website. Four predicted genes, including *CD97*,*Hmbg1,Cadm1* and *Srsf1* with high expression level in GT1-7 cells were assumed as the candidate target of miR-505-3p. Dual luciferase reporter assay in HEK293 cells were performed to test the inhibitory effect of miR-505-3p by binding to their 3’UTR. The results indicated that the other three genes except for CADM1 could be the target genes of miR-505-3p (S5 Fig A, B). Integrated with their functional annotation, *Srsf1* was selected preferentially for further investigation. Given that SF2 was proved to be a regulator of mTOR signaling, we analyzed the phosphorylated status of S6K in pGT1-7 cells, but the amount of p-S6K made no difference between GT1-7 and pGT1-7 cells (S5 Fig C).

The accumulation of SF2 protein was less in pGT1-7 cells than in GT1-7 cells (Fig. 7A). Knocking down *Srsf1* expression in GT1-7 by shRNA inhibited the expression of *Kiss1* and *Gnrh* simultaneously. When the expression of *Srsf1* was rescued in pGT1-7 cells with *Srsf1*steadily overexpressed, the expression level of *Kiss1* and *Gnrh* increased (Fig. 7B). These results indicated that miR-505-3p may inhibit the expression of puberty-related genes through *Srsf1*.

**Fig 7:**
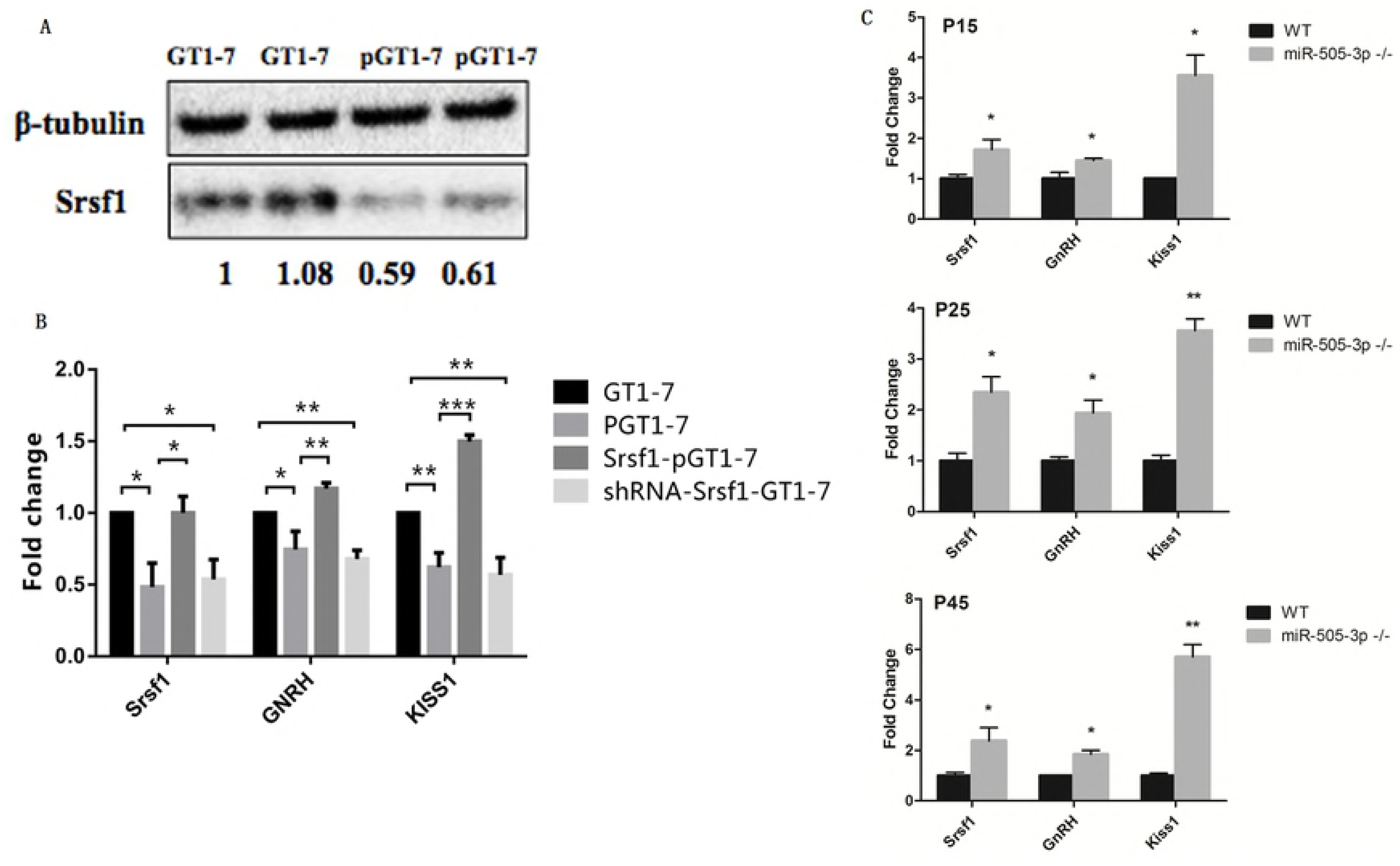
miR-505-3p and its target gene *Srsf1* play a role in the puberty onset regulation. A: The translation of Srsf1 was inhibited by miR-505-3p in pGT1-7 cell line. B: The expression level of puberty related genes in pGT1-7 after Srsf1 overexpression. Bars are means and vertical bars represent SEM (* P<0.05, ** P<0.01, *** P<0.001, ns no statistical significance). C: Srsf1, Kiss1 and GnRH expression level in the hypothalamus of miR-505-3p KO mice at PND15, PND25, PND45

Moreover, *Srsf1, Kiss1* and *GnRH* in hypothalamus of miR-505-3p knockout mice and wild mice at different postnatal days were also detected, and the result showed knockout mice had higher expression levels (Fig. 7C).

### RIP-seq results showed SF2 mainly bound to RP mRNAs in GT1-7 cell line

RNA immunoprecipitation sequencing (RIP-seq) was used to identify RNAs which bound to SF2 in GT1-7 cell line. The SF2 protein and its bounding RNAs were pulled down by SF2 specific antibody, and the RNA component was sequenced by next generation sequencing. We pulled down IgG protein and its bounding RNAs as one control sample, and the RNA extracted from GT1-7 was another control sample. The RNA-seq data of the SF2 pull down sample was compare to the two control data separately, and the two comparing results were highly consistent, SF2 protein mainly bound ribosome related mRNAs in the GT1-7 cell (Table 2).

**Table 2.**
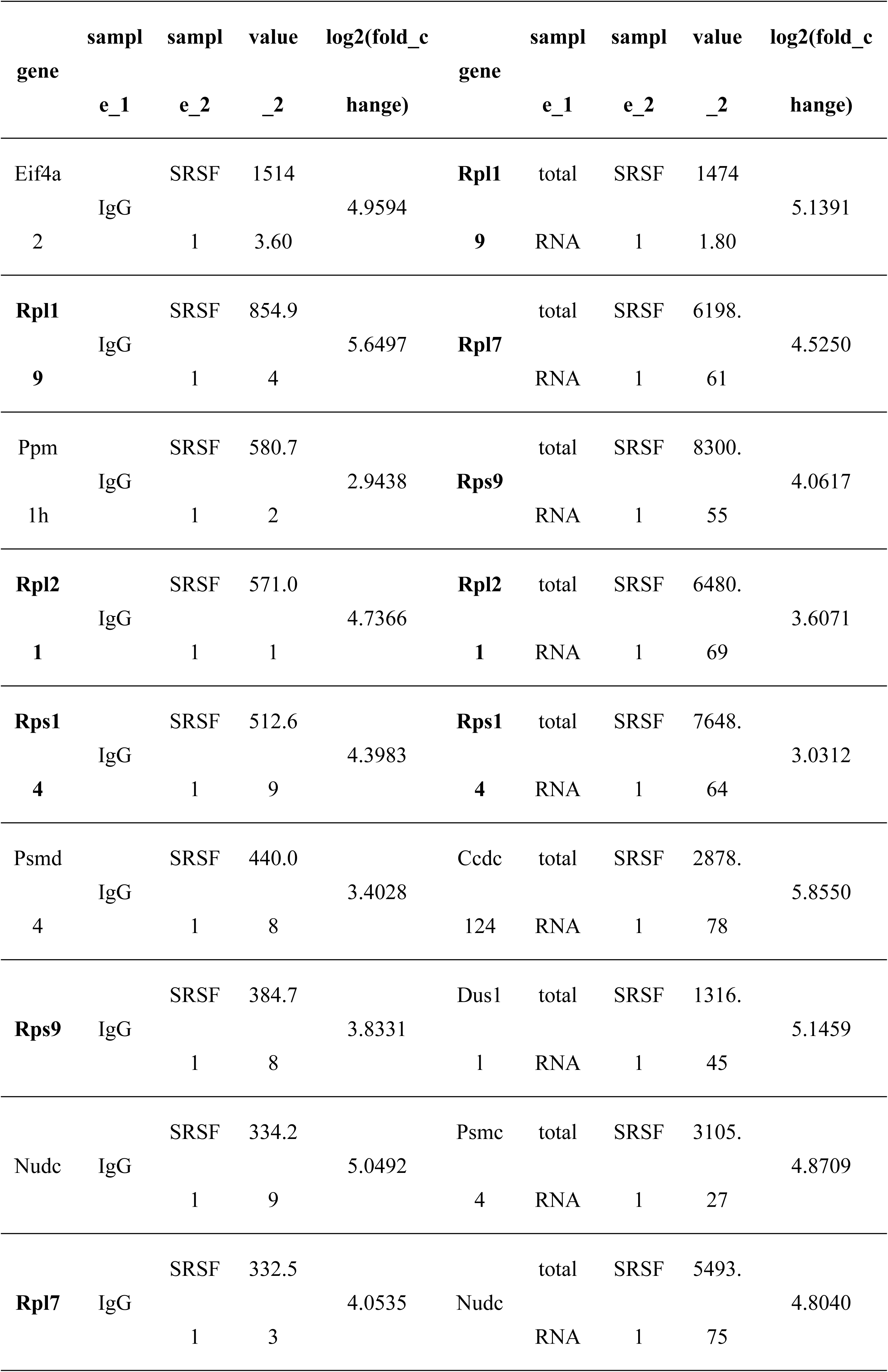
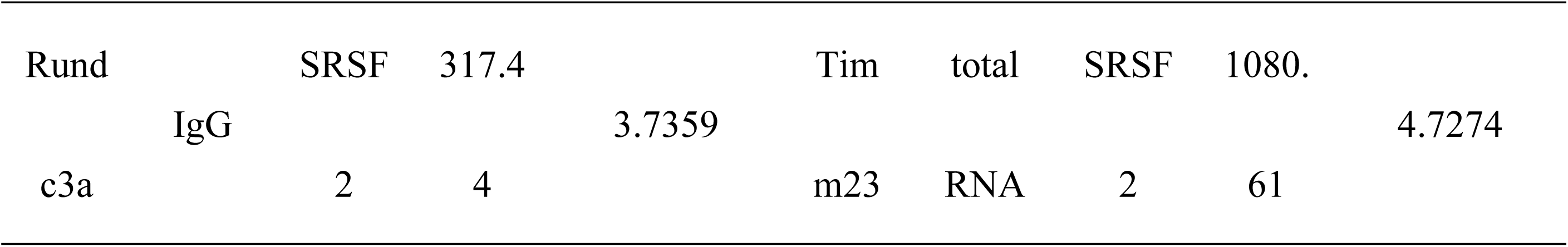
Top 10 gene list of SF2 pull down sample RNA-seq data compare to two control samples’ data separately

We performed the KEGG analysis on the transcriptome data compared pGT1-7 with GT1-7 cell. The result showed ribosome and ribosome biogenesis in eukaryotes pathway were both affected in pGT1-7 cell line (Fig. 8).

**Fig 8.**
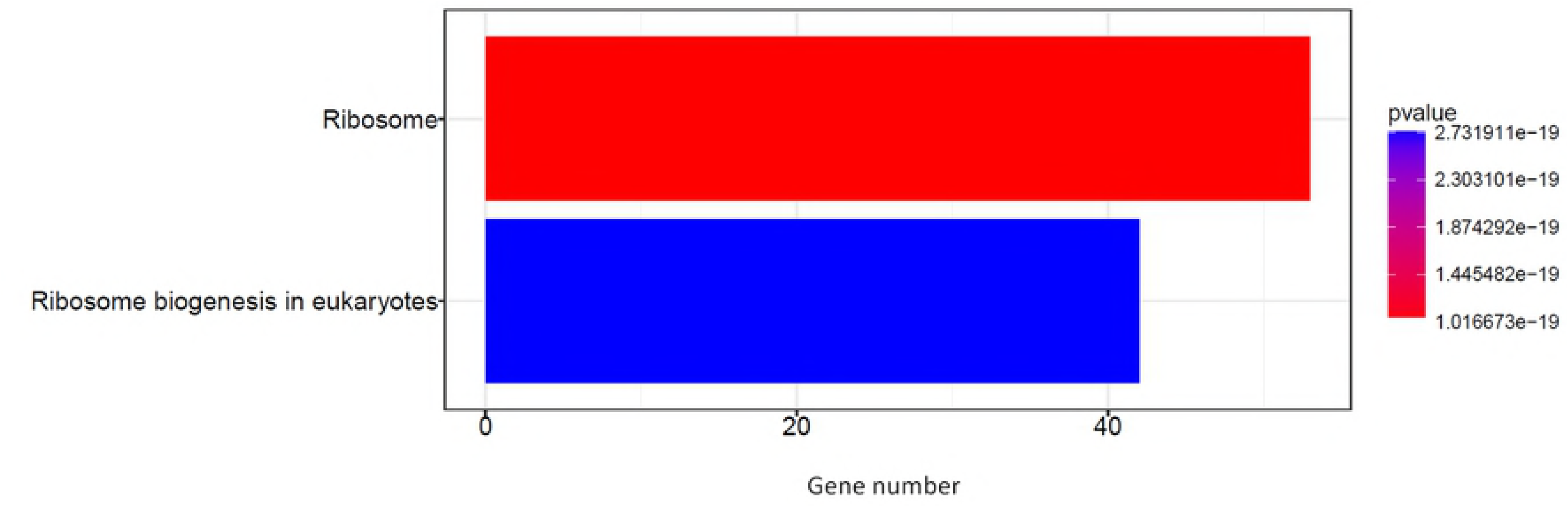
Pathway analysis of differentially expressed genes between GT1-7 and pGT1-7 cell.

## Discussion

Genetic strategies, such as positional cloning and association study, can be powerful tools to reveal the functional genes underlying the complex traits [10, 21–23]. We took the advantage of mouse models to make interval-specific congenic strains (ISCS) between C3H/He and C57/BL6 mice to finemap the QTLs related to the puberty onset and then reveal the candidate gene according to the sequence variation, expression differentiation and functional connection. To our knowledge, miR-505-3p is the first microRNA which was found by positional cloning as a puberty regulator in female mice.

Considerable progress has been made to decipher the molecular foundation of the pubertal timing [24–26]. Especially some studies revealed the potential epigenetic regulatory role of miRNAs and lncRNAs in female puberty[27]. Okabe et al indicate that the knockout of miR-200b and miR-429 can induce reduced fertility in female mice^[^28^]^.Zhu et al show the variation in Lin28/let7 pathway can change the body size and onset of puberty in mice[29]. Furthermore, miR-132, miR--9 and miR-145 are also involved in the pubertal timing owing to their potential regulatory role of c-myc and Lin28^[30,31]^. In 2016, Messina et al. reported a microRNA switch that controled the rise of hypothalamus GnRH production before puberty in mice [32]. They provided direct evidence that the increasing expression of miR-200/429 and miR-155 in GnRH neurons during the infantile period lifted the repressive control of GnRH expression, by repressing ZEB1 and CEBPB, respectively [32]. These results suggested the existence of a multilayered and interconnected array of miRNAs and their target genes that control the puberty timing. Our results shows the overexpression of miR-505-3p in the hypothalamus can cause the delay of puberty timing and reduced fertility in female mice, and the results from miR-505-3p knockout mice supported the deduction, which could be a new member of this regulating array.

We constructed a miR-505-3p ectopic expression female mouse model by lentiviral delivery. Lentiviral vector as a versatile tool has been accessible to transfect post-mitotic neurons and induces substantial, long-term transgene expression^[^33^-^35^]^. In our experiment, the miR-505-3p overexpression in the mouse hypothalamus directed by lentivirus reached the peak (about 25 times as the control) on the tenth day after injection and continued for at least 40 days (though declined to 8 times of the control), which provided an ideal window to investigate the puberty onset and reproductive actions of the female mice affected by the microRNA. Besides reverse transcription and real-time PCR, we performed in situ hybridization for miR-505 detection using improved LNA-probes with high specificity and sensitivity. The expression pattern of hypothalamic miR-505-3p detected by in situ hybridization can be used to further elucidate the relationship between miR-505-3p function and the location of miR-505-3p in the hypothalamus. Actually we have observed some connections between the injecting site and the phenotypic distinction.

Finding out the target gene of miR-505-3p underlying the puberty onset in female mice can help to tell the regulation mechanism of this complex trait. In our previous work, we identified one of the important functions of miR-505-3p as that it was a crucial regulator for axonal elongation and branching through modulating autophagy in neurons [36]. Atg12 was testified to be the key target gene of miR-505-3p in this process, as its protein product ATG12 (autophagy-related 12) was an essential component of the autophagy machinery during the initiation and expansion steps of autophagosome formation. However, we did not find the influence of miR-505-3p overexpression on the amount of ATG12 in GT1-7 cells, either in the mRNA expression profiling data or qRT-PCR detection, which could be due to the different cell types we used. Autophagy is more active in neurons than in cultured cells [37, 38], and the autophagy-related proteins in neurons could be not as sensitive to the microRNA modulation as in the cultured GT1-7 cells. We found that when cultured in serum reduced medium, the target gene of miR-505-3p became more sensitive to microRNA regulation in GT1-7 cells [39], which implied that the decreased velocity of the protein synthesis under malnourished condition can help the fine tuning of microRNAs on their targets. *Srsf1* is the first experimentally validated target gene of miR-505-3p, and it was also a regulator of mTOR pathway, which was reported to participate in the VO modulation in rats [20]. However, in miR-505-3p overexpressing pGT1-7 cells, the amount of the marker of the activated mTOR signaling phosphorylated S6K was not affected, which implied that the inhibition of the *Srsf1* expression by miR-505-3p did not influence the mTOR signaling in GT1-7 cells.

*Srsf1* is the archetype member of the SR (Serine an Argenine rich protein) family of splicing regulators and has multiple functions in the cell nucleus and cytoplasm. Moreover, *Srsf1* is also a proto-oncogene in cell malignant transformation [40, 41]. SF2 could bind to ribosomal proteins and help in mRNA stability, transport, intracellular localization, and translation [42]. Ribosomal proteins play independent key roles in the regulation of apoptosis, multidrug resistance and carcinogenesis [43,44], RPL22 can inhibit Lin28B, RPS7 regulates PI3K/AKT and MAPK [45]. Our RIP results showed that SF2 bind to the mRNA of ribosomal proteins preferentially which had not been reported before. The biological significance of this selectivity has not been clarified though, it encourages us to assume that miR-505-3p may affect the independent roles of ribosomal proteins by inhibiting SF2 in GT1-7 cells, and the output of these series effects is the downregulation of puberty related genes. The potential relationship between RP and the puberty development needs further elucidation. The function of miR-505-3p as a puberty regulator in female mice has been revealed by cell and animal model, but the underling mechanism still requires further researches.

## Materials and methods

### Fine mapping of the puberty-related QTL on Chromosome X in mice

In our previous work, a significant puberty-related ChrX QTL of 2.5 cM was found by genome scanning, in a panel of 10 modified interval-specific congenic strains (mISCSs) in C57/BL6 and C3H/He mice [10]. In this study, the mice of the strain carrying the QTL were backcrossed with C3H mice. The resulting male F2 mice holding at least one recombination at the special interval were chosen and then backcrossed with female C3H to obtain N2 generation. The female N2 continued to mate with male C3H to generate N3 generation. N3 male mice holding only one recombination at the target interval were selected and continued to backcross with female C3H to generate N4. N4 mice siblings were crossed until N7 generation. At last, the age at VO of all female mice of N7 progenies was recorded. All modified ISCSs were verified by genotyping for the genetic markers on chromosome X. All animal procedures were approved by the Animal Ethics Committee of Donghua University, and all experiments were conducted in strict accordance with the National Institutes of Health Guide for the Use of Laboratory Animals.

### DNA Extraction and Sequencing

DNA Extraction from the tail tip was performed using a DNA Extraction Kit (Tiangen, China), followed by the instructions provided by the manufacturer. All gene sequence data were from NCBI database. And the primers were designed by Primer3 (http://frodo.wi.mit.edu/primer3/) and synthesized by Shanghai Sangon Biotech Ltd. PCR was performed in 15 μL reaction mix containing 3 μL DNA template, 200 nM of each PCR primer, 0.25 mM of each dNTP, 3 nM MgCl2, 1.5 μL 10×PCR Buffer and 1 U Hot start DNA Polymerase (Tiangen, China) covered by mineral oil. The reaction was carried out at 95°C for 15 min, followed by 95°C 30 sec, 55°C 0.5 min, 72°C 1 min, 35 cycles. The purified PCR products were sequenced by 3730XL sequencing instrument (ABI, USA).

### SNP database querying between C3 and B6 mice

All SNPs between C3 and B6 mice in the QTL region (rs13483770∼rs299055848) on chromosome X were queried from Mouse Genome Project in Sanger Database (http://www.sanger.ac.uk/science/data/mouse-genomes-project).

### qRT-PCR

Total RNA from hypothalamus or 10^6^ cells was isolated using Trizol reagent (Invitrogen, USA) in line with the manufacturer’s protocol. 1ug of total RNA was reverse-transcribed with oligo-dT (for mRNA detection) or microRNA specific primer (for microRNA detection) by RevertAid First strand cDNA synthesis Kit (ThermoFisher, USA). Real-time PCR was performed using SuperReal PreMix Plus (TIANGEN,Beijing, China) on 7500 Real-time PCR System (Applied Biosystems) and normalized to *Actb* (for mRNA) or miR-16 (for microRNA). All reactions were run in triplicate and included no template control for each gene. All PCR primers were listed in S3 Table.

### Generation of hypothalamic miR-505 overexpressing mice

C57/BL6 (B6) inbred mice were purchased from Shanghai SLAC Laboratory Animals Co.LTD (Shanghai,China). The miR-505-3p carrying lentivirus delivery was performed on female mice at postnatal 12-15d weighted at 5.2-5.6 g. After 1% sodium pentobarbital anaesthesia, mice were fixed on stereotaxic apparatus (STOELTING, USA). According to Paxions et al^[21]^, The hypothalamic stereotaxic coordinates were carried out relative to the bregma (anteroposterior [AP]= -1.53mm and lateral [L]=±0.24mm) and to the skull (dorsoventral[DV]=-5.3mm). The injection of lentiviral preparation (2.83E+08 IU/ml, 900 nl per sample) was at the rate of 30 μL/min. Saline were also injected to the female mice at the same condition to be used as controls. After injection, the mice were kept at 37°C till recovered from anesthesia. Afterwards, the mice were put back to their mothers for further investigation.

### Cryostat section

Mice were under 1% sodium pentobarbital anesthesia and perfused by 0.9% saline. When the color of liver and lung become pale, saline was replaced by 4% paraformaldehyde to fix the tissues. Then the brains and ovaries were taken out and kept in paraformaldehyde solution at 4 °C overnight. The fixed brains and ovaries were dehydrated by glucose solution and cut into sections using Cryotome (Thermo,USA). The thickness of brain and ovary sections was 15μm and 20μm, respectively. The sections were attached to slides which were pretreated with potassium dichromate and embedded in polylysine. Slides were stored at -80 °C until objected to FISH and HE staining, respectively.

### In situ hybridizations

Brain sections were taken out from -80 °C and dried at room temperature for at least 10 min. The sections were fixed by 4% paraformaldehyde and pretreated withj Triton x -100 and Proteinase K to improve the tissue permeability. After pre-hybridization, sections were hybridization with diluted digoxigenin (DIG)-labeled LNA probes for miR-505-3p, (5’-AGAAAACCAGCAAGTGTTGACG-3’) at 55°C overnight. Then the sections were washed and incubated with Horseradish Peroxidase(HRP) conjugated anti-DIG antibodies(Abcam,Shanghai,China) at 4.°C overnight. The fluorescent signals were amplified using TSA^TM^Cyanine 5 Systerm (PerkinElmer,USA) and detected by inverted phase contrast fluorescence microscope (Olympus, Japan).

### Measurement of pubertal events and fertility analysis

The animals were housed under the specific pathogen free (SPF) standard conditions with a 12 h light/dark cycle and adequate water and food. Injected female pups were caged with their mothers again and weighed every two days. After weaned at postnatal 21d, the mice were examined daily for the vaginal opening. They were mated with fertile male mice at the age of 8 weeks at the ratio of two to one. Female mice with drastic body weight gaining were assumed to be pregnant and were separated from male mice into a single cage. The interval time between mating and delivery was recorded. Female mice that failed to be pregnant for more than one month were regarded as infertile. The rate of infertility in female mice and the death rate of newborns were calculated. Mice with vacant virion or saline injected were used as control.

### Hormone detection

Serum LH and FSH were extracted by mouse orbital blood sampling. The detection of LH or FSH was performed using Mouse LH ELISA KIT or Mouse FSH ELISA KIT(Elabscience,Wuhan,China) according to manufacture’s procedures, respectively. The standard curve was plotted by CurveExpert 1.4 software, which was also used to calculate the concentration of LH and FSH referring to the standard curve.

### Histological analysis

Ovarian sections were taken out from -80 °C and dried for at least 10 minutes. Then the sections were dehydrated by ethanol and stained by hematoxylin and eosin (HE). The morphology of ovaries was detected by light microscope (Olympus, Japan).

### Generation of miR-505 knockout mice

The miR-505 knockout mice were generated in CRISPR/Cas9 technology by The Animal Facility of Indtitute of Neuroscience, Chinese Academy of Sciences. Female B6 mice were superovulated using pregnant mare serum gonadotropin (PMSG) and were injected with human chorionic gonadotropin (HCG) after 48 h. The superovulated female mice were mated with B6 stud males, and fertilized eggs were collected from their oviducts after 36 hours. Two sgRNAs were designed to guide Cas9 targeting the pre-miR-505 locus (S1 Fig). A mixture of transcribed Cas9 mRNA and sgRNA was microinjected into zygotes of B6 mouse. The injected zygotes were transferred into the uterus of pseudopregnant female mice immediately after injection. The pregnant mice were housed in standard cages on a 12 h light/dark cycle with *ad libitum* access to food and water till delivery.

The pups were taken by tail clips and their DNAs were extracted for sequencing. Pups with large deletions in pre-mir-505 region were reserved as founders. The generated male and female founders were mated with wild-type B6 mice at the age of 40 days. The heterozygote F1 mice were self-crossed to generate double knockout (DKO), single knockout (SKO) and wild type mice for further puberty events study. The resulting female mice were genotyped by PCR detection with the primers, forward: 5’-AAACCAGCAAGTGTTGACGC, reverse: 5’-CCCTGTTTGTCACTTGCAGA, The lengths of the PCR products were 120bp for wild type, 103bp ` 120bp and 97bp ` 120bp for heterozygous, and 97bp and 103bp for homozygous knockout mice.

### Construction of miR-505-3p stable over-expressed and *Srsf1* knockdown GT1-7 cell line

The double strand DNA sequence of miR-505-3p was cloned into the pLenti6.3/V5-DEST lentiviral expression vector (Invitrogen, USA). The target sequence of shRNA for *Srsf1* (5’-GCCCAGAAGTCCAAGTTAT-3’) and non-specific shRNA (5’-TTCTCCGAACGTGTCACGT-3’) were synthesized and cloned into the PLKD-CMV-R&PR-U6-shRNA lentiviral expression vector. The resulting lentivirus vectors were were packaged into virions by Obio Technology Co. LTD (Shanghai, China). The virions were used to infect GT1-7 cells. Cells were seeded 100000 cells per well in twenty-four-well plate and cultured in DMEM medium with 10% FBS (Gibco, USA). After 4 days of transfection, cells were selected under antibiotics according to the vector’s resistances for 14 days. After 10 days of expansion without antibiotics, the surviving cells were harvested for further investigation. GT1-7 cells with miR-505-3p overexpression were named as pGT1-7.

We used GeneChip Mouse Transcriptome Assay 1.0 (Affymetrix, USA) to get the transcriptome data of pGT1-7, and GT1-7 as the control. Differential expressed genes were used to perform KEGG analysis based on Kyoto Encyclopedia of Genes and Genomes database.

### Cell culture and RIP-seq

GT1-7 cells are immortalized cell lines derived from mouse hypothalamic neurosecretory neurons which provide an ideal model system for study of that regulate reproduction as their similarity to GnRH neurons, such as the synthesis, processing, and pulsatile secretion of GnRH (gonadotropin-releasing hormone)[47]. GT1-7 cells applied in this study were kindly provided by Professor Xiaoying Li of Shanghai Clinical Center for Endocrine and Metabolic Diseases, Shanghai Jiaotong University suggested by Dr. Pamela Mellon at University of California, San Diego. We performed RNA immunological precipitation (RIP) assays followed by high-throughput sequencing in SF2 pulldown of GT1-7 cell, IgG pulldown and total RNA as the control. Briefly, after cell lysis, 10% of sample removed for input stored at -80°C until ready to perform RNA purification. The input sample is used to calculate RIP yields, and may also be used to evaluate the quality of the RNA. Add RNA binding SF2 protein-specific antibody or negative control IgG to supernatant. Incubate with rotation at 4°C for overnight. Add washed protein A+G Agarose to each IP sample and incubate with rotation at 4 °C for 2 hours, then wash off unbound material and remove supernatants. RNAs bound to RNA-binding protein were purified with Trizol and sequenced by NGS.

## Acknowledgments

We appreciate Professor Xiaoying Li in Shanghai Clinical Center for Endocrine and Metabolic Diseases, Shanghai Jiaotong University for providing GT1-7 cells. We also thank Yu Li and Li Chen for help doing some experiments, Fuyi Xu for the assistance of data analysis, and Xiaoning Li for the assistance of manuscript writing.

